# Generation of guard cell RNA-seq transcriptomes during progressive drought and recovery using an adapted INTACT protocol for *Arabidopsis thaliana* shoot tissue

**DOI:** 10.1101/2021.04.15.439991

**Authors:** Anna van Weringh, Asher Pasha, Eddi Esteban, Paul J. Gamueda, Nicholas J. Provart

## Abstract

Drought is an important environmental stress that limits crop production. Guard cells (GC) act to control the rate of water loss. To better understand how GCs change their gene expression during a progressive drought we generated guard cell-specific RNA-seq transcriptomes during mild, moderate, and severe drought stress. We additionally sampled re-watered plants that had experienced severe drought stress. These transcriptomes were generated using the INTACT system to capture the RNA from GC nuclei. We optimized the INTACT protocol for *Arabidopsis thaliana* leaf tissue, incorporating fixation to preserve RNA during nuclear isolation. To be able to identify gene expression changes unique to GCs, we additionally generated transcriptomes from all cell types, using a 35S viral promoter to capture the nuclei of all cell types in leaves. These data sets highlight shared and unique gene expression changes between GCs and the bulk leaf tissue. The timing of gene expression changes is different between GCs and other cell types: we found that only GCs had detectable gene expression changes at the earliest drought time point. The drought responsive GC and leaf RNA-seq transcriptomes are available in the Arabidopsis ePlant at the Bio-Analytic Resource for Plant Biology website.

## Introduction

Guard cells (GC) help plants survive periods of drought by reducing or halting water loss from transpiration (Nilson and Assmann, 2006). With a predicted increase in the incidence of drought from climate change models, there is growing importance to understand how plants are able to perceive, respond to and mitigate stress (Cominelli et al., 2009; Yusa et al., 2015). The identification of genetic players can guide the identification of mutants or the engineering of transgenics with improved stress tolerance. Transcriptomic analysis captures snapshots of the broad gene expression patterns that may be driving cellular responses and previous studies have aimed to understand the genes that regulate GC behaviour. Early GC transcriptomes aimed to characterize gene expression changes in response to the stress hormone abscisic acid (ABA) that is able to induce rapid stomatal closure at high concentrations (Yang et al., 2008; Leonhardt et al., 2004; Wang et al., 2011). Further investigations profiled the GC response to sucrose (Bates et al., 2012) and reduced humidity (Bauer et al., 2013). Despite the interest of GCs in drought, only one GC transcriptome has characterized the response to drought treatment, and this was a prolonged exposure to moderate drought levels (Prasch et al., 2015).

GC transcriptomes have previously been generated using a few different techniques to isolate the cell type. GCs have strengthened cell walls and can tolerate higher mechanical forces from blending (Prasch et al., 2015; Bauer et al., 2013) and can handle a longer digestion time with cell wall digesting enzymes than other cell types (Yang et al., 2008; Wang et al., 2011; Leonhardt et al., 2004). Necessarily, these approaches cannot be used to directly compare to any other cell type, but they have allowed for comparisons of GCs of different genotypes and under different treatments. Protoplasting, however, has been noted for having an influence on the RNA pool, with a minimum of a few hours of digestion at non-zero temperatures being required (Bates et al., 2012). While new transcription can be blocked by transcriptional inhibitors, RNA degradation has been seen to occur in a non-random way in GC samples with short versus long digestion times (Obulareddy et al., 2013).

There are methods for cell type-specific transcriptome analysis that would allow for comparison between different cell types, including Fluorescence Activated Cell Sorting (FACS), Isolation of Nuclei TAgged in specific Cell Types (INTACT; Deal and Henikoff, 2011) and Laser Capture Microdissection (LCM). FACS and INTACT require transgenic plants, limiting their quick application to mutants of interest. The INTACT method has the benefit of allowing for the isolation of nuclear RNA from frozen tissue thereby obviating the need for protoplasting enzymes, but FACS still requires a protoplasting step to release individual cells. LCM requires processing time and skill to manually dissect tissue types of interest. Of these, only LCM has been used to generate a GC transcriptome (Bates et al., 2012), although protoplasting without FACS has been used too (Yang et al., 2008).

We generated RNA-seq data sets profiling gene expression changes in GCs through a physiologically relevant progressive drought and after recovery. We sampled plants experiencing mild, moderate, and severe drought. We chose the INTACT method to capture our cell type-specific responses and generated bulk leaf tissue controls for all time points and treatments. This is the first study that applies the INTACT method to adult *Arabidopsis thaliana* leaf tissue. As we observed RNA degradation during nuclear processing following the published INTACT protocol, we substantially adapted the protocol to include fixation during tissue collection and the subsequent release of RNA after fixation. Differential gene expression analysis identified 4837 differentially expressed genes (DEGs) in GCs and 6551 in leaves across these time points, with a combined total of 7075 unique DEGs identified across all tissues and time points. These data sets identified that GC drought responsive gene expression changes are unique and shared. Notably, GCs alone responded to the mild drought condition.

## Methods

### Generation of INTACT transgenic plants

A 3.3kb KAT1 promoter (Mustroph et al., 2009) was amplified from *A. thaliana* Col-0 genomic DNA and the 35S promoter sequence was amplified from the pEGAD plasmid, adding Gateway sequence tags for BP cloning. Promoters were cloned into the INTACT backbone plasmid (Kanamycin resistance, obtained from Brady lab, UC Davis) using Gateway enzymes (Invitrogen). Plasmids were transformed into competent *Agrobacterium tumefaciens* and a floral dip was performed on *A. thaliana* Col-0 plants to generate T1 resistant plants (Clough and Bent, 1998). Single insertion mutants with a 75% inheritance pattern of the INTACT backbone in the T2 generation and the subsequent homozygous progeny (100% resistance) were identified for both transgenic lines. GFP expression patterns were assessed (Leica SP5) in T2 and T3 seedlings to ensure these lines had the desired cell type-specificity.

### Progressive drought and re-watering treatment

KAT1:INTACT and 35S:INTACT seeds were sown in 9cm pots, filled with an equal weight of Sunshine Mix #1 soil (Sun Gro Horticulture). Pots were stratified for 3 days at 4°C in the dark, then moved to growth chambers with a 9-hour daylight program with a growth temperature of 21°C in the light and 18°C in the dark and a leaf height light intensity of 120 μmol m^-2^ s^-1^. Seedlings were thinned to one seedling per pot after germination. Drought treatment began with a final day of watering when plants were 4.5 weeks old and all pots were watered until the same weight range was reached for all pots, about 90% soil water content (SWC). Drought treatment plants were weighed daily to track SWC, each pot had a unique identifying number to track the drought progression of each individual pot and control plants were watered regularly to maintain a SWC above 80%. Plants and age-matched well-watered controls were sampled when drought treated plants reached 60%, 40% and 20% SWC. On the day of 20% sampling, some 20% SWC pots were re-watered and sampled 24 hours into their recovery. All samples were collected 4.5 hours after the start of the 9-hour daylight cycle. Two progressive drought experiments were performed: one drought was performed collecting both KAT1:INTACT and 35S:INTACT plants, the second drought was performed to collect additional replicates for KAT1:INTACT plants. In total around 200 35S:INTACT and 700 KAT1:INTACT plants were grown and collected for these experiments.

At time of sampling, whole shoots were cut at the base and placed into DEPC treated jars (wide mouth 250mL Bernardin jars) with 100-200 mL of room temperature 4% paraformaldehyde (PFA) in PBS, sufficient volume to fully submerge the plant tissue. Plants were placed under vacuum for 1.5 hours to allow the fixative to fully penetrate the tissue. To stop the fixation, plants were washed once with ice cold 125 mM glycine PBS solution and once with PBS. Jars were placed on ice during all washes and until tissue was ground. With all liquid poured off, fixed plants were ground to a fine powder in liquid nitrogen and stored at −80°C until nuclear isolation. All solutions were DEPC treated or RNAse free and mortar and pestles were treated with RNAseAWAY (Sigma).

### Processing of nuclei, RNA extractions and RNA sequencing

Nuclei were isolated following the INTACT protocol (Deal and Henikoff, 2011), with minor modifications. Nuclear Purification Buffer (NPB): 20 mM Tris-HCl pH 7.5 (Invitrogen), 1% PVP, 40 mM NaCl, 90 mM KCl, 2 mM EDTA pH8, 0.5 mM EGTA pH 8.0, 0.2 mM Spermine, 0.5 mM Spermidine, 0.5x Protease inhibitors (cOmplete, Roche) made in RNAse free water (Ultrapure, Invitrogen). A 90 second 500 × *g* spin was added to clear the supernatants before pelleting nuclei at 1200 × *g* for 7 min (Reynoso et al., 2017). Isolated nuclei were resuspended in proteinase K buffer (PKB): 10 mM Tris-HCl pH 8.0 (Invitrogen), 0.5% SDS (Promega 10% SDS solution), 0.2 M NaCl, 50 mM EDTA pH 8.0. To release RNA from fixation, nuclei were incubated with Proteinase K (Thermo Scientific) for 15 min at 55°C, incubated at 70°C for 30 min with a 40 mM concentration of Tris-HCl pH 8.0 (Evers et al., 2011; Slane and Bayer, 2017) then transferred directly into Trizol. Trizol:chloroform supernatants were mixed 1:1 with 70% ethanol and transferred to RNeasy micro columns for RNA extraction. RNA was treated with Turbo DNase (Thermo Fisher) and cleaned using ethanol precipitation, with sodium acetate and glycogen (Thermo Scientific). RNA quality and quantity were assessed using Pico Bioanalyzer Chips (Agilent) and Qubit RNA HS Assay kits (Thermo Fisher). RNA libraries were prepared using TruSeq Stranded Total Plant RNA with Ribo-Zero Plant (Illumina), where rRNA depletion precedes library preparation. 30 million 75bp reads were generated for each sample using the NextSeq500 Desktop Sequencer v2 (Illumina). All samples were sequenced in triplicate, with the exception of KAT1:INTACT at 40% SWC, which was sequenced in duplicate and KAT1:INTACT at 20% SWC was sequenced in quadruplicate. We attempted to get triplicates for KAT1:INTACT at 40% SWC, but some samples failed to generate libraries despite having similar RNA quality and quantity^1^.

### Mapping, data pre-processing, and differential expression analysis

RNA-seq reads were mapped to the TAIR10 genome (NCBI Refseq GCA_000001735.1) with Hisat2 (Pertea et al., 2016) using the read strand flag, all other settings were default. Reads that mapped to nuclear chromosomes were submitted to Stringtie (Pertea et al., 2016) to produce gene counts for protein coding transcripts, using known splice sites from the Araport11 genome annotation. One protein coding gene (AT2G01021) was identified and removed from subsequent analysis as a source of bias that we did not want to influence differential gene expression analysis or between sample normalization, since this gene was highly expressed. This gene is located near to the rRNA locus on Chromosome 2. Sequencing from our nuclear RNA showed high levels of external transcribed spacers (ETS) and internal transcribed spacers (ITS), which are part of the polycistronic rRNA. There was some technical variation in how well each sample depleted rRNA genes, and fluctuations in the gene AT2G01021 increased along with rRNA reads.

Differential expression analysis was performed on raw read counts per gene in edgeR (Robinson et al., 2009). Analysis was done to test for differential expression between treatments of the same genotype (drought and re-watering) against their age matched well-watered controls. Comparisons between genotypes was done testing for differences between all well-watered samples of each genotype. To correct for batch effects from KAT1:INTACT samples being collected across two drought experiments, the drought was specified in the model in edgeR when testing for differential expression along with the sequencing lane (Leek et al., 2010).

### Gene count normalization, clustering, and GO term enrichment

Gene lengths were calculated as the average length of all isoforms using gene positions noted in the Araport11 gff3 annotation file. Transcripts Per Million (TPM) counts were calculated in edgeR to normalize counts between samples. K-means clustering was performed in R using base packages. The total within-cluster sum of squares for different numbers of clusters were calculated to avoid overfitting the data with too many clusters. Heatmaps were generated in R using the package pheatmap2. GO term enrichment analysis for lists of differentially expressed genes was performed using AgriGO v2 (Tian et al., 2017) Singular Enrichment Analysis.

## Results

### INTACT system for RNA profiling in *A. thaliana* leaf tissue

To profile GCs and bulk tissue using the same experimental approach, we established the INTACT system for RNA profiling in Arabidopsis leaf tissue. INTACT allows for extraction of cell type specific nuclei for RNA or chromatin profiling (Deal and Henikoff, 2011). We generated *A. thaliana* Col-0 plants harboring KAT1:INTACT and 35S:INTACT constructs. We validated the cell type specificity of these plant lines. GFP expression in KAT1:INTACT plants was restricted to GCs and 35S:INTACT GFP was observed in all examined leaf cell types (**Figure 1**, Supplemental Figure 1). Initial attempts to isolate nuclei from *Arabdiopsis* leaf tissue according to the standard INTACT protocol resulted in severe RNA degradation (bulk of RNA <200nt, data not shown). The INTACT protocol was then adapted to include paraformaldehyde fixation of live leaf tissue, in particular whole shoots, before freezing and storage at −80°C. RNA was then released from fixation before RNA extraction (see Materials and Methods for details). This modification to the protocol allowed us to obtain RNA of sufficient yield and quality for library preparation from Arabidopsis green tissue (**Figure 2**), the inclusion of fixation protected RNA from degradation during nuclear processing. Importantly, the cell type-specificity of the biotinylated tag is compatible with formaldehyde fixation, as formaldehyde fixation is included in the INTACT protocol for profiling chromatin prior to nuclear isolation (Deal and Henikoff, 2010).

**Figure 1.**
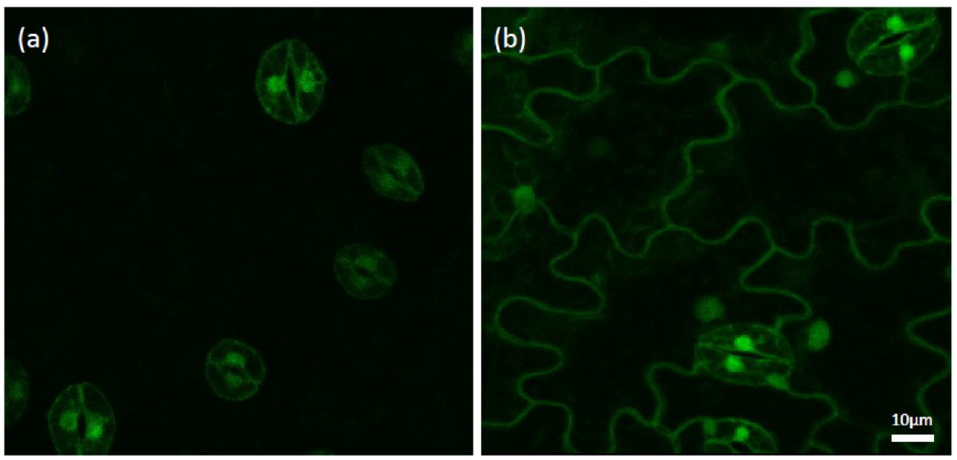
GFP expression in abaxial epidermis of true leaves of 2-week-old seedlings (a) KAT1:INTACT and (b) 35S:INTACT.

**Figure 2.**
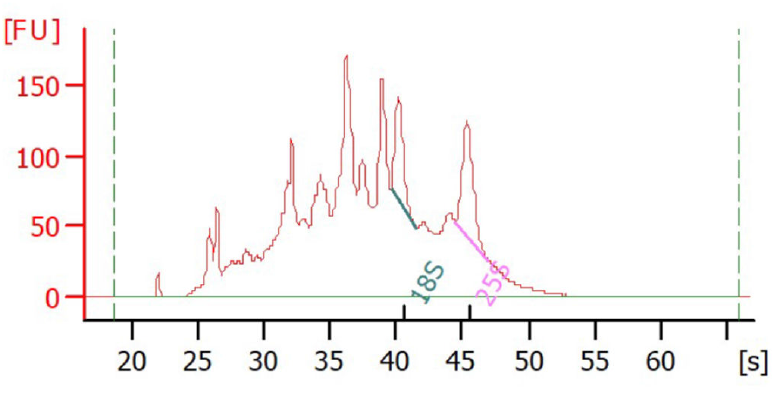
Representative Bioanalyzer trace of RNA obtained from 5-week-old KAT1:INTACT plants using modified INTACT protocol from paraformaldehyde fixed tissue.

### Guard cell and whole shoot RNA-seq transcriptomes during progressive drought and recovery

KAT1 and 35S:INTACT plants were subjected to a physiologically relevant progressive drought treatment. Pot weights were assessed daily for drought treated plants to track the onset of drought severity. We sampled plants through a drought that set in over 2 weeks, capturing plants experiencing mild (60% SWC), moderate (40% SWC) and severe (20% SWC) drought stress (**Figure 3**). Age-matched well-watered controls were also sampled at all drought time points. To profile the response to drought recovery, plants that reached 20% SWC were re-watered and sampled 24 hours later. All samples were collected in the middle of the daylight cycle, 4.5 hours into a 9-hour day. This captures GCs when they would be fully open in the diel cycle under non-stressed conditions (Dodd et al., 2004).

**Figure 3.**
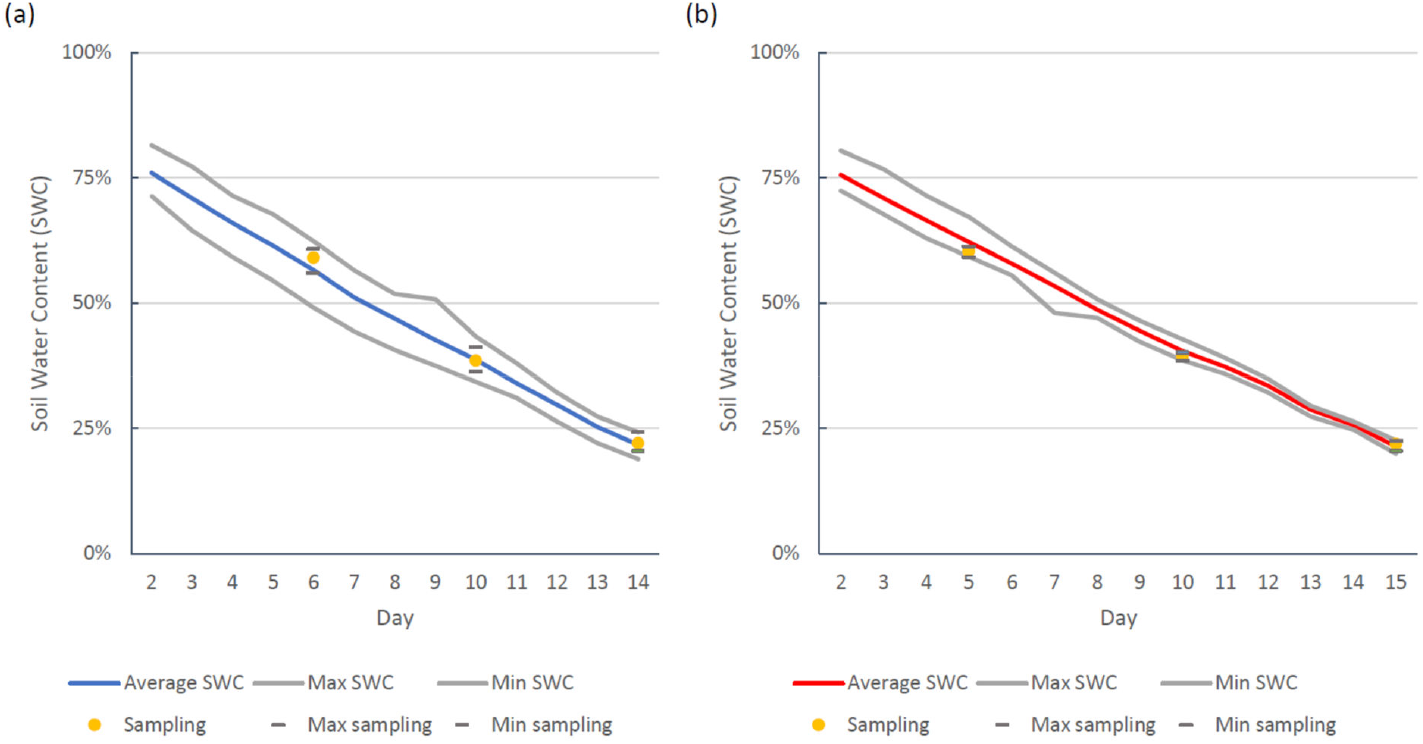
Soil water content (SWC) measurements from plants sampled in the progressive drought experiment. The average, maximum and minimum SWC for sampled pots is displayed on the day that sampling occurred. SWC measurements include plants from two drought experiments for (a) KAT1:INTACT and one drought experiment for (b) 35S:INTACT.

RNA was processed using our modified INTACT protocol and 30 million RNA-seq reads were generated for each RNA sample. All samples were sequenced in triplicate except for KAT1:INTACT at 40%, where only two samples were able to generate libraries (see Material and Methods for discussion of library failures). An additional KAT1:INTACT at 20% was sequenced, making a total of four replicates for this time point.

### Mapping, expression level quantification, and quality control

Reads were mapped to the *A. thaliana* genome, and gene counts for known nuclear protein coding gene transcripts from Araport11 were quantified. These samples contained a clear biological signal from both genotype and treatment, principal component analysis using raw RNA-seq counts for all genes separated the two genotypes on the first axis and treatment along the second (Supplemental Figure 2a). Early drought treatments and well-watered samples were not as clearly separated than the more severe drought and re-watering samples, indicating that these early drought treatments had a smaller impact on the transcriptome. We additionally noted that 35S:INTACT replicates clustered more separately than KAT1:INTACT. We also note age related clustering for the well-watered time points, despite only being 4-5 days apart. This supports the need for age matched controls in analyzing stress-responsive gene expression through a time course.

KAT1:INTACT plants required more plants for each replicate, to obtain sufficient RNA for these rare cell types. To collect sufficient samples and RNA, some KAT1:INTACT plants at the 20% time point (all conditions) were collected from two separate drought experiments. We noted a drought effect, where KAT1:INTACT samples from Drought 1 and Drought 2 were separated in our principal component analysis (Supplemental Figure 2b). The drought was consequently included in the model for differential expression analysis to control for larger effects that the drought had on gene expression in these replicates. Having samples from two drought treatments may reduce the number of gene expression changes detected, but these gene expression changes would represent changes shared between two slightly different drought treatments.

### Differential Gene Expression from Droughted GCs and Shoots using INTACT Nuclear Profiling

We tested for significant differential expression for all treatment conditions against age matched well-watered controls in edgeR. We additionally tested for significant differential expression comparing re-watered to 20% drought samples, as severely droughted plants were those subjected to re-watering treatment. The number of differentially expressed genes (DEGs) for each comparison is summarized in **Table 1**. A response to mild drought stress was detected in GCs but very few changes were detected in whole shoots. 95 DEGs exhibited increased transcript abundance and 75 DEGs decreased in abundance in GCs while only 2 DEGs with increased expression levels were detected in whole shoots. The number of DEGs increased with drought severity for both GC and whole shoots, with thousands of DEGs detected in severe drought stress and after re-watering. In total across these time points, DEG analysis identified 4,837 stress-responsive genes in GCs, of which 3,973 occurred during drought stress. Whole shoots had a higher total with 6,551 stress-responsive genes, of which 5,460 occurred during drought stress. Across the two genotypes there were 9,618 unique genes with significant gene expression differences in any treatment, and 7,075 of those unique genes were altered during drought.

**Table 1.** Counts of significant differentially expressed genes (DEG) detected by edgeR using a false discovery rate of 0.1 and percent of DEGs with a magnitude greater than 2-fold change for all comparisons. “Up” denotes genes with increased expression levels relative to the controls, and “Down” denotes genes with decreased expression levels relative to the controls. Well-watered samples were combined (for 9 replicates) to test for DEGs between genotypes (GC vs Shoot). All drought treatments (60%-20% Soil Water Content, SWC) were tested for significant differential expression against their age matched well-watered (ww) control, re-watered samples were tested against their age-matched well-watered controls and against the 20% SWC treatment. All samples were sequenced in triplicate except for GC 40% SWC drought treatment, which was in duplicate, and GC 20% SWC treated samples, which were in quadruplicate.

|  |  | Up<br>> 1.4 fold change | Down<br>> 1.4 fold change | Up<br>> 2 fold change (%) | Down<br>> 2 fold change (%) |
| --- | --- | --- | --- | --- | --- |
| Genotypes | GC vs Shoot | 2273 | 1862 | 40% | 24% |
| GC | 60% SWC | 95 | 75 | 28% | 19% |
|  | 40% SWC | 277 | 105 | 39% | 46% |
|  | 20% SWC | 1407 | 2230 | 23% | 30% |
|  | Re-water vs 20% | 2106 | 1652 | 26% | 26% |
|  | Re-water vs ww | 685 | 1099 | 23% | 26% |
| Whole shoot | 60% SWC | 2 | 0 | 100% | *n/a* |
|  | 40% SWC | 326 | 550 | 15% | 21% |
|  | 20% SWC | 2018 | 3255 | 26% | 39% |
|  | Re-water vs 20% | 3320 | 1840 | 40% | 28% |
|  | Re-water vs ww | 1102 | 1023 | 23% | 15% |

Gene expression changes in re-watering encompassed a mix of recovery-specific and lingering drought-responsive DEGs. In GCs, 423 (64%) of DEGs whose transcript levels increased and 441 (38%) of DEGs whose transcript levels decreased in the re-watered vs well-watered samples are not DEGs in drought. In whole shoot it is 628 (56%) of DEGs with increased abundance and 463 (45%) with decreased abundance. Indeed, nearly 80% of drought responsive DEGs return to well-watered levels within this 24-hour recovery period with only 10% unchanged by watering, except genes with decreased levels of expression in GCs that retain nearly 20% of genes at stress-responsive levels (Supplemental Table 1).

### Guard cells and whole shoots have shared and unique patterns of differential gene expression

As show by k-means clustering of 9,618 expression profiles of genes differentially expressed at any timepoint or in any genotype (KAT1:INTACT or 35S:INTACT) in **Figure 4**, there are patterns of increases in expression at late drought timepoints that are both shared in guard cells and leaves (like those in cluster A) or that are unique to guard cells (cluster B). Also, the direction of responses is sometimes similar in guard cells and leaves, but the magnitude can vary, as in cluster H, during re-watering. These data may be explored for any gene with the BAR’s ePlant tool at http://bar.utoronto.ca/eplant/ (Waese et al., 2017). The expression patterns for *At4g05100* (*AtMYB74*), a transcription factor shown to be induced under salt stress (Xu et al., 2015) is depicted using this tool in **Figure 5**. This representation shows the importance of having age-matched controls, as this gene appears to increase in expression in the guard cells of well-watered plants over the two-week course of the experiment.

**Figure 4.**
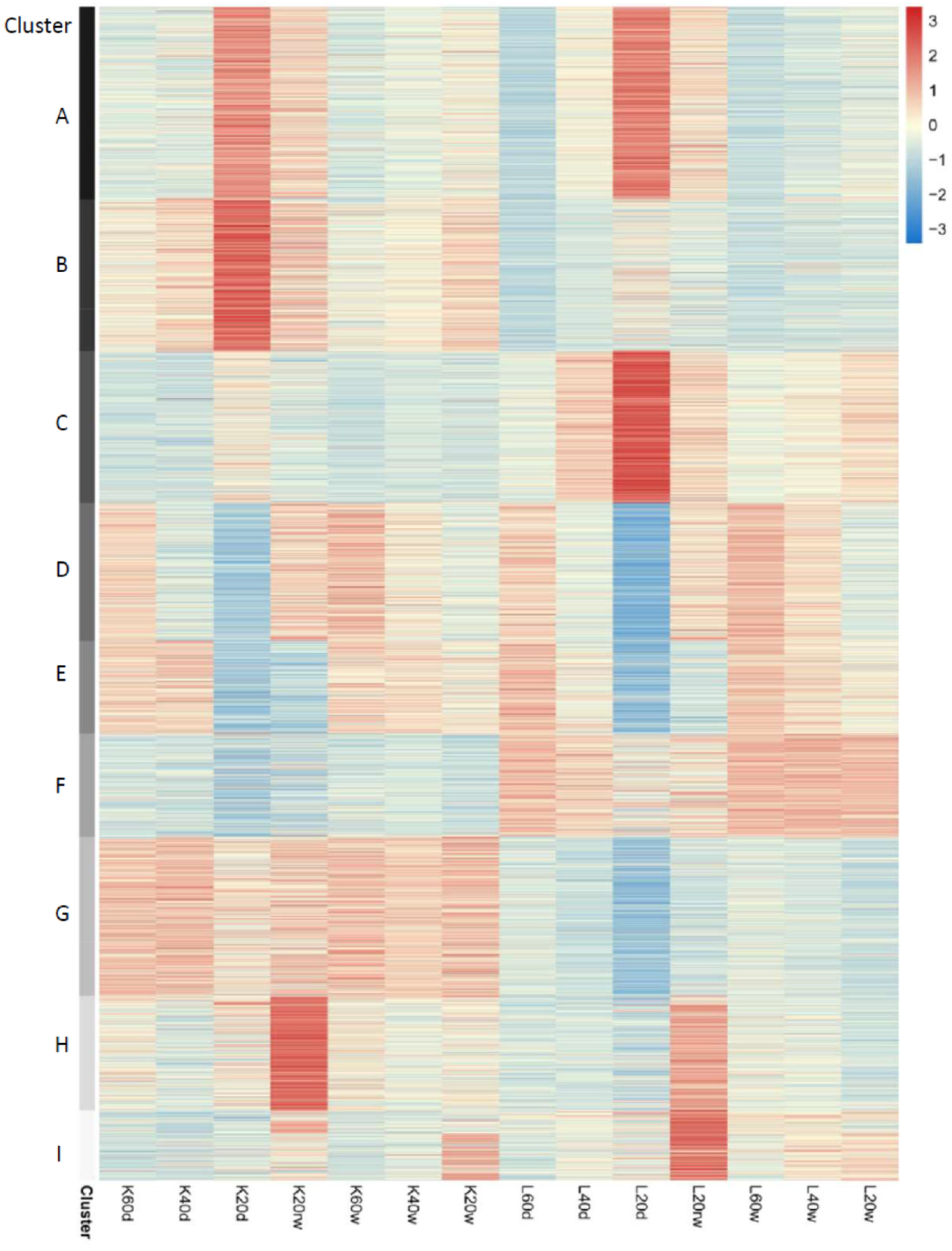
Heatmap of treatment responsive DEGs from both GCs and whole leaves. K-means clustering with 9 clusters was used to group 9,618 genes by unique and shared expression patterns in samples. Samples are labelled to show the genotype, time point, and treatment: *K* represents GCs sampled using KAT1:INTACT, *L* represents whole shoots sampled using 35S:INTACT; *60,40* or *20* represents the time point referring to the %SWC of the paired drought sample; *w* is for well-watered, *d* is for droughted, and *rw* is for re-watered samples.

**Figure 5.**
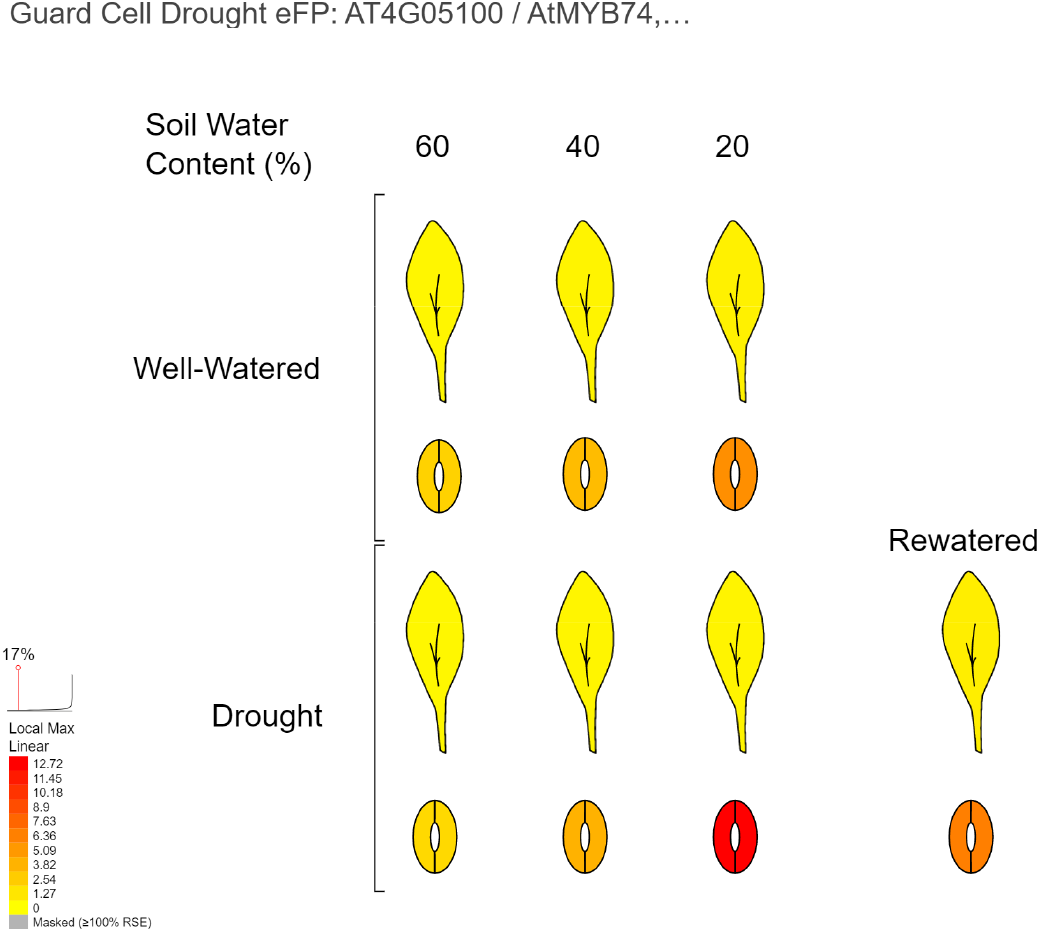
The expression patterns for *At4g05100* (*AtMYB74*), a Cluster B gene from the previous figure, in the Guard Cell Drought eFP view of the BAR’s Arabidopsis ePlant by Waese et al. (2017), shared under a CC BY licence. This depiction shows the importance of having age-matched controls, as this gene appears to increase in expression in the guard cells of well-watered plants over the two-week course of the experiment, even in the absence of drought. Expression values are TPM.

### Novel stress-responsive gene expression changes in these data sets

To understand how much of the GC drought responsive gene expression observed in our data set overlapped with previously published transcriptomes, we compared our DEG lists with those from 6 other studies (**Table 2**). Drought responses are a mixture of ABA-dependent and independent responses and GC-specific data sets have not previously looked at a progressive drought. We therefore chose three GC ABA responsive transcriptomes (Leonhardt et al., 2004; Wang et al., 2011; Bauer et al., 2013), two whole leaf progressive drought treatment transcriptomes (Bechtold et al., 2016; Harb et al., 2010) and a GC-specific data set profiling plants exposed to moderate drought over 5 days treatment (Prasch et al., 2015). For comparison we also examined the overlaps for drought responsive DEGs that we identified in whole shoots.

**Table 2.** Proportion of DEGs identified in our analysis shared in DEG lists from 6 chosen publications or in a merged set of all DEGs for the 6 GC ABA or drought responsive data sets combined. Genes were considered shared by data sets if the direction of the change was the same, e.g. a DEG exhibiting increases in expression in our data sets would be a shared gene if was also found to increase in expression levels in a given ABA/drought responsive data set, but it would not be a shared gene if that gene were found to be decreased in expression in a given ABA/drought responsive data sets from the 6 chosen publications (or combination) examined here. Directionality “up” or “down” refers to increased transcript abundance, or decreased transcript abundance, respectively, as compared to the appropriate control. SWC is soil water content.

|  | Treatment | Directionality | GC transcriptomes * | GC transcriptome ^ | Leaf transcriptome ’ | Leaf transcriptome ’ | Leaf transcriptome ’’ | 6 transcriptomes * ^ ’ ’’ |
| --- | --- | --- | --- | --- | --- | --- | --- | --- |
|  |  |  | ABA responsive | moderate drought acclimation | moderate drought | progressive drought | progressive drought | ABA and drought |
| GC | 60% SWC | up | 25% | 1% | 18% | 35% | 6% | 48% |
|  |  | down | 15% | 4% | 25% | 21% | 13% | 59% |
|  | 40% SWC | up | 22% | 4% | 13% | 29% | 8% | 43% |
|  |  | down | 4% | 1% | 10% | 13% | 6% | 26% |
|  | 20% SWC | up | 16% | 8% | 15% | 28% | 23% | 47% |
|  |  | down | 8% | 8% | 12% | 31% | 14% | 53% |
| Whole shoot | 60% SWC | up | 0% | 0% | 0% | 0% | 0% | 0% |
|  |  | down | n/a | n/a | n/a | n/a | n/a | n/a |
|  | 40% SWC | up | 10% | 2% | 7% | 17% | 13% | 30% |
|  |  | down | 7% | 7% | 13% | 17% | 11% | 40% |
|  | 20% SWC | up | 13% | 3% | 10% | 24% | 14% | 37% |
|  |  | down | 10% | 4% | 10% | 29% | 9% | 46% |

\* Leonhardt et al. (2004), Wang et al. (2011), Bauer et al. (2013)
^ Prasch et al. (2015)
‘ Harb et al. (2010)
“ Bechtold et al. (2016)

Overall, all time points seemed to have a combination of known ABA responsive genes in GCs, known drought responsive genes and gene expression changes unidentified in these six data sets. In GCs there were between 26 and 59% of DEGs shared with these other six data sets. The highest overlaps with the GC data sets were observed with the ABA responsive data sets and the progressive drought experiment from Harb et al. (2010). There was generally more overlap with ABA data sets in our GC data for genes increased in expression, and more so for early responding genes (at 60% SWC). With the Harb et al. (2010) moderate and progressive drought time points, there was overlap with genes found to both increase and decrease in expression levels, and the progressive (more severe drought time point) time point did have a slightly higher overlap with our 20% SWC time point than the moderate drought time point. Generally, there were more overlaps seen with the GC data sets than the whole shoot data sets, even though the drought treatments were sampling whole leaf.

Little overlap was observed with the Prasch et al. (2015) data set, which might have been expected to exhibit a high amount of concordance due to the shared cell type and drought, however a sustained drought treatment can be considered more of an acclimation experiment, as plants are able to adapt within a few days to a sustained moderate drought, including a diminution of ABA responses within 2 days of moderate drought (Harb et al., 2010). Additionally, we sampled at midday while Prasch et al. (2015) sampled at the end of the daylight cycle, and stress-responsive gene expression changes are known to be strongly influenced by the time of day (Wilkins et al., 2010).

### GO term enrichment analysis

We submitted individual DEG lists for genes with increased (“up”) transcript abundance or for genes with decreased (“down”) transcript abundance as compared to age- and tissue-matched well-watered controls to AgriGOv2. A composite figure describing significantly enriched GO terms for all soil water content samples and tissues was generated and the results of this analysis may be seen in **Figure 6**. It is apparent that certain changes – in carbohydrate metabolism genes, for example – are occurring specifically in guard cells (see red highlighted areas in Figure 6), although there are also categories that are enriched across several comparisons.

**Figure 6.**
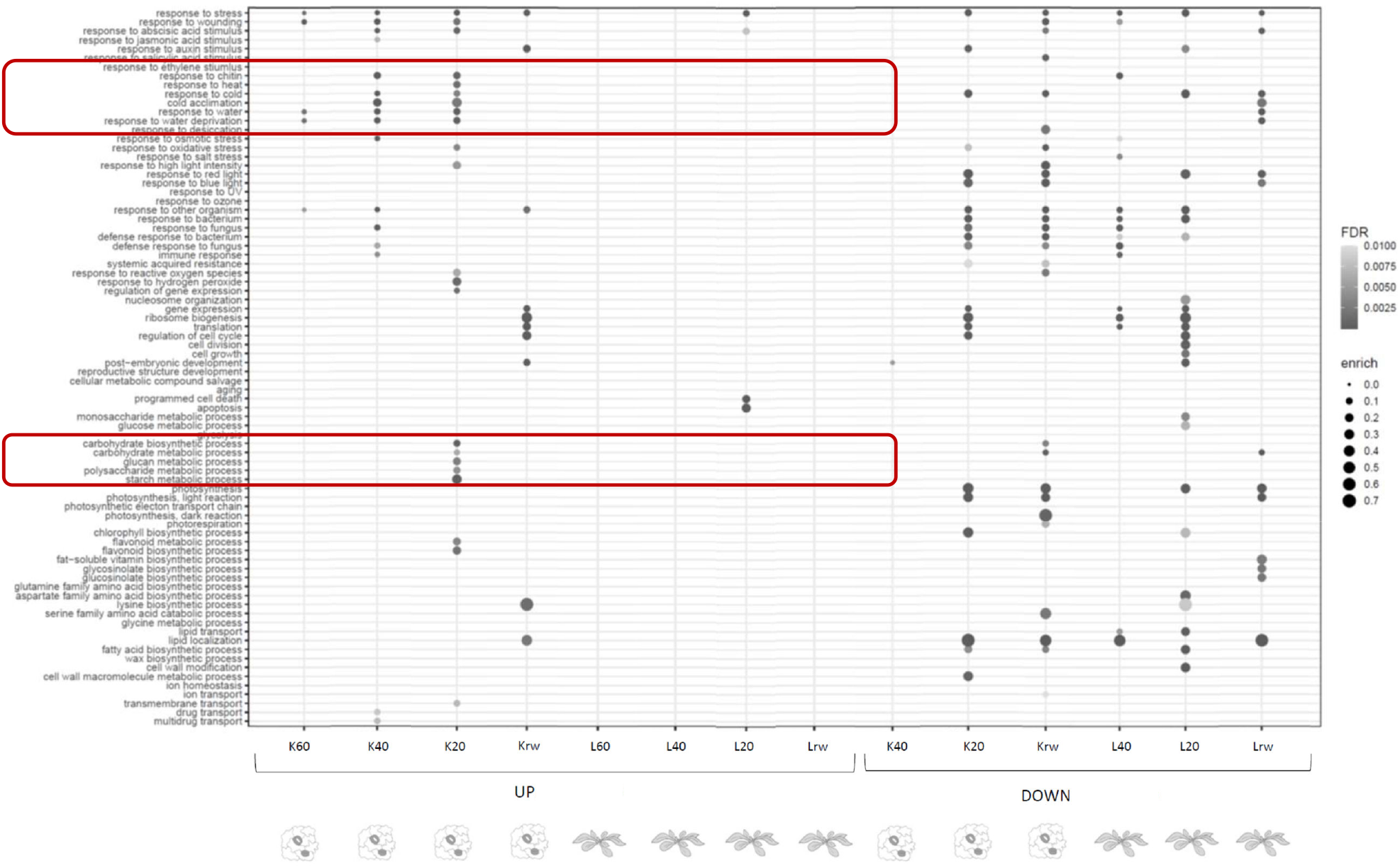
Significantly enriched GO terms for DEGs from GCs and whole leaf samples for all treatments vs their matched well-watered control. “Up” and “Down” DEGs (i.e., those genes exhibiting increases or decreases, respectively, in transcript abundance compared to their age- and tissue-matched well-watered controls) were submitted for GO term enrichment analysis. Samples are labelled to show the genotype time point and comparison: *K* represents GCs sampled using KAT1:INTACT, *L* represents whole shoots sampled using 35S:INTACT; *60, 40* or *20* represents the time point referring to the % SWC (soil water content) of the paired drought sample. Red highlighted areas denote terms enriched specifically for GC “up” DEGs as compared to whole leaf samples.

## Discussion

We present the first GC-specific transcriptomic experiment profiling a a novel RNA-seq data set that captures gene expression changes in guard cells in response to a gradually developing progressive drought. The method to isolate guard cell RNA is improved and involves a fixation step to preserve the quality of RNA prior to nuclei isolation. By using a guard cell-specific promoter and one that drives ubiquitous expression in bulk leaf tissue, we were able to compare whole leaf and guard cell-specific drought responses, which permitted us to identify shared and unique responses in each. Understanding what is unique to guard cells will help guide us in terms of engineering of guard cell behaviour in response to drought.

There have been many experimental approaches to study drought, from a “quick and dirty” approach – for example the Kilian et al. (2007) drought treatment involved growing plants on rafts and removing these rafts from the hydroponic growth medium and then drying the plants in an air stream until 10% of the fresh weight was lost, versus more a physiological approach (e.g. Bechtold et al., 2016). We would stress the importance of studying a progressive drought because this reflects the slow onset of drought experienced in nature. Harb et al.’s (2010) and Bechtold et al.’s (2016) progressive drought captured how changes in gene expression change with increasing stress in bulk leaf tissue, but did not capture responses at the level of guard cells. The concept of stress memory has been explored in Arabidopsis plants by removing them from soil, air-drying them to “train” them, then rehydrating them at high humidity, and repeating the cycle several times (Ding et al., 2013). This study highlighted the path dependency of stress responses, at least in the bulk leaf tissue used in these studies. No studies, however, have generated guard cell transcriptomes from a progressive drought. The only drought responsive data set to date is from Prasch et al. (2015), which held plants at a moderate drought levels for five days, a level that was considered tolerable by the plants.

In our experiment, we observed that guard cells – but not the bulk leaf control – responded at the level of gene expression to mild drought stress (i.e. at 60% and 40% SWC), as shown by the red highlighted areas in Figure 6 (other GO categories of specific responses also exist, but are not highlighted). Guard cells are early targets for drought signals, for example by the drought-induced hormone ABA, because of ABA importers present in guard cells (Kang et al., 2010), although it is unknown whether there is any ABA being produced with such mild changes in water availability.

The modified INTACT protocol we developed for whole shoots could be used to extract nuclear RNA for any cell type in that tissue using the same protocol and thus extends the range of application to many different areas of research outside of roots, where the INTACT system is well established. Of course, the applications require the use of transgenic plants expressing the nuclear targeting fusion tag in the desired cell type. Yields are manageble although the scale of experiment would depend on the cell type: many more plants were needed for every guard cell sample in our experiments than for the ubiquitously-expressed nuclear targeting fusion tag driven by the 35S promoter. Nevertheless, given the ever smaller amounts of RNA required for RNA-seq, exploring cell types that comprise less than 1-2% of the cell types (as is the case for guard cells) is feasible, even if many plants are required. That samples can be fixed, frozen and processed later is a large advantage of the INTACT method, especially for environmental stress perturbation experiments. Quick collection – by collecting fixed, whole shoots – combined with nuclear RNA extraction would, for example, be ideal for dissection of transcriptional responses requiring very frequent sampling. The fixation step also permits the same samples to be used for chromatin accessibility studies (Deal and Henikoff, 2010).

Whether the continued differential expression of some drought responsive genes is associated with a behavioural change in guard cells is unknown, but the differential expression of some genes in guard cells from rewatered plants as compared to those from plants at 20% SWC would suggest this to be the case. Maintaining stress-responsive gene expression patterns after the resolution of the stress could imply a “hedging” strategy and could affect guard cell behaviour under new drought conditions. It has been previously noted in potato plants that stomata (formed by guard cell pairs) take a few days to return to normal conductance levels after re-watering after a drought experiment (Kopka et al., 1997). It is possible that the genes that remain at 20% SWC stress levels could dampen guard cell responsiveness after re-watering. As our experiment was interrogating nuclear RNA, we hypothesize that these stress-responsive genes are still being newly transcribed despite the return to normal water levels. Perhaps this means that these genes do indeed play a role, and that these are genes responding to drought signals that do not return to pre-drought levels as quickly because they offer some continued benefit to plant survival and recovery from stress, or exposure to subsequent stress.

The Bechtold et al. (2016) drought kinetics were very similar to ours yet we saw little overlap in terms of differentially-expressed gene sets for our bulk leaf samples – Bechtold et al. also examined whole leaf response. This could be reflective of the type of RNA we collected (nuclear instead of mRNA), the profiling platform we used (RNA-seq versus CATMA microarrays), the type of tissue we harvested (we used whole shoots while Bechtold et al. used Leaf 7, or growth conditions. That there was little overlap with the Prasch et al. (2015) guard cell data set is more easily explained by the time of day influence when samples were taken – Prasch et al. collected at end of diel cycle while our samples were collected midday. Last, under the mild drought acclimation experiment conducted by Harb et al. (2010) prolonged ABA production stops, and there’s also a change in the growth rate, thus making it difficult to directly compare these data sets. Nevertheless, we believe that our data set and modified INTACT approach represent extremely useful additions to the literature.

We have made the guard cell drought response RNA-seq data set available in the BAR’s ePlant under the “Guard Cell Drough eFP” in the “Tissue & Experiment eFP viewers” module. Future work in our laboratory will aim to further understand what roles these guard cell-specific, drough-responsive genes play.

### Contributions, Acknowledgements and Data Access

AvW developed the modified INTACT protocol, conducted the drought experiment, performed RNA isolations, mapped RNA-seq data, undertook the DEG analyses, and created all figures except Figure 5, which was generated by ePlant. PJG summarized expression levels and AP databased the summarized expression levels on the BAR. EE created the ePlant “Guard Cell Drought eFP” input image. AvW and NJP wrote the manuscript. All authors have approved the manuscript. NJP is grateful for funding from the National Sciences and Engineering Research Council of Canada, through the Discovery Grants program, and from Genome Canada / Ontario Genomics (OGI-071) for the original ePlant framework. The expression data may be freely explored at http://bar.utoronto.ca/eplant/ – enter a gene and once it has loaded, select the “Guard Cell Drough eFP” in the “Tissue & Experiment eFP viewers” module.

## Supplemental Materials

**Supplemental Figure 1.**
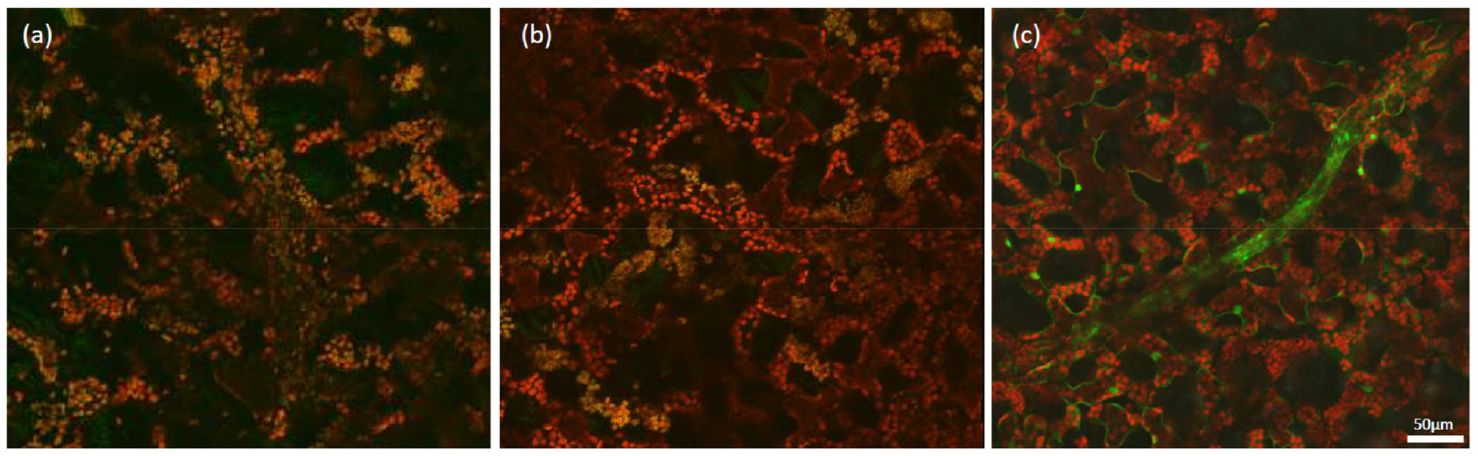
GFP expression in mesophyll and vasculature, revealed by peeling away abaxial epidermis of 5-week-old leaves of Arabidopsis plants (a) Col-0 untransformed wild-type, (b) KAT1:INTACT, and (c) 35S:INTACT. GFP signal is overlayed with chlorophyll autofluorescence, shown in red.

**Supplemental Figure 2.**
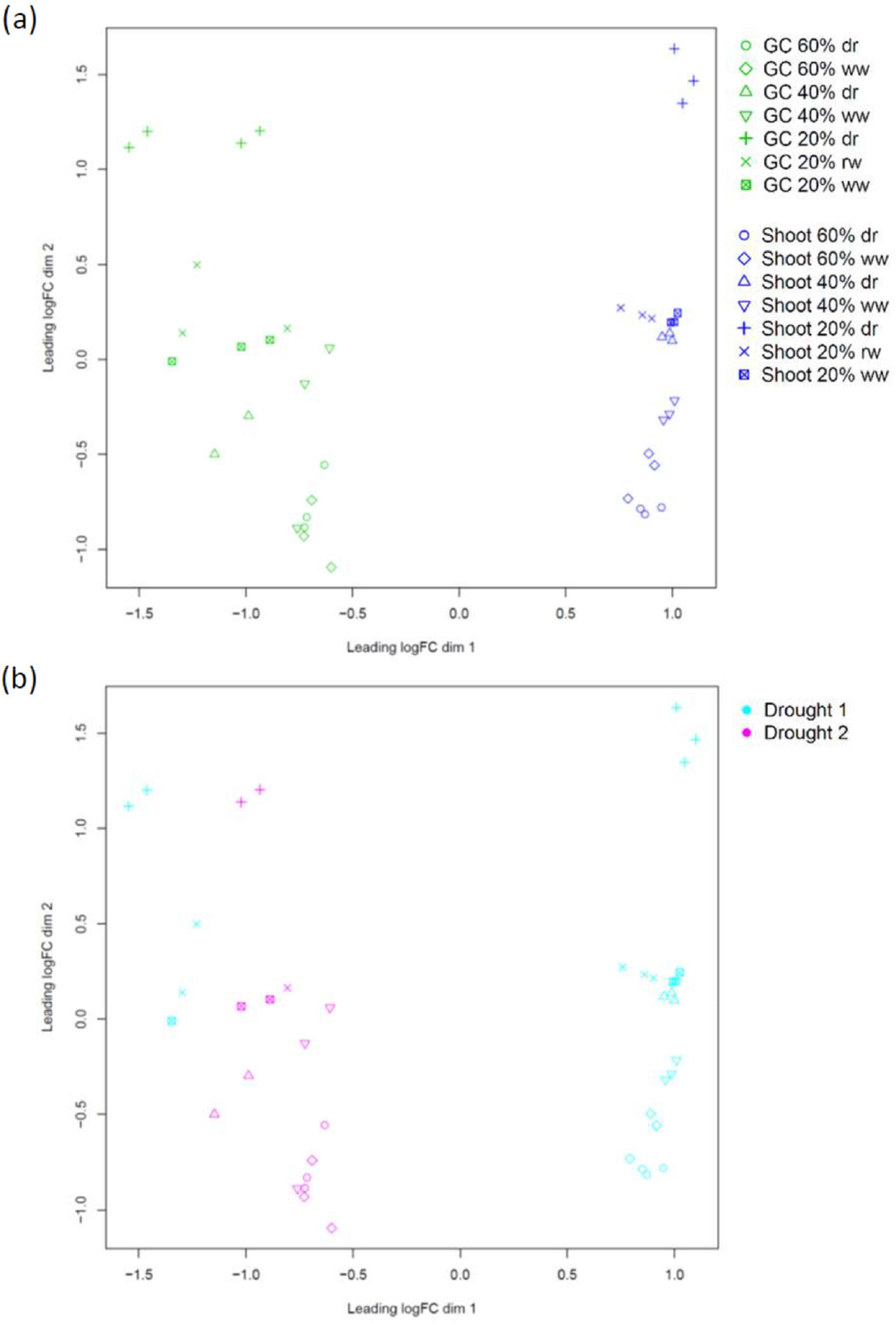
Principle component analysis of raw counts for (a) replicates of samples. Samples collected from Drought 1 and Drought 2 are highlighted in (b).

**Supplemental Table 1.** Subcategorization of 20% DEGs based on whether DEG between rewatered (rw) vs 20% soil water content (SWC) or rewatered vs well-watered (ww), and in which direction (up or down, indicating increased transcript abundance or decreased transcript abundance, respectively) for these comparisons. *More so*: DEG at rw vs 20% SWC drought, and in the same direction at 20% SWC vs ww. *Stays*: Not DEG in rw vs 20% SWC drought, but DEG in rw vs ww in same direction. *On the way back*: DEG in the same direction as 20% SWC drought vs ww and rw vs ww, but opposite direction (and DEG) in rw vs 20% SWC drought. *Back to normal*: not DEG in rw vs ww. *Reverses*: DEG at rw vs ww and in opposite direction from DE at 20% SWC drought vs ww. GC refers to guard cellspecific data set.

| Genotype | Timepoint | Trend at rw | Up | Down |
| --- | --- | --- | --- | --- |
| GC | 20% SWC | More so | 5 (0.4%) | 8 (0.4%) |
|  |  | Stays | 143 (10.2%) | 492 (22.1%) |
|  |  | On the way back | 18 (1.3%) | 73 (3.3%) |
|  |  | Back to normal | 1193 (84.8%) | 1587 (71.2%) |
|  |  | Reverses | 48 (3.4%) | 70 (3.1%) |
| Shoot | 20% SWC | More so | 5 (0.2%) | 5 (0.2%) |
|  |  | Stays | 238 (11.8%) | 360 (11.1%) |
|  |  | On the way back | 29 (1.4%) | 125 (3.8%) |
|  |  | Back to normal | 1686 (83.5%) | 2602 (79.9%) |
|  |  | Reverses | 60 (3.0%) | 163 (5.0%) |

## Footnotes

1 A few KAT1:INTACT samples from various conditions and time points not able to be used to prepare libraries. No failures were observed for 35S:INTACT plants. Generally, the RNA quantities for KAT1:INTACT samples were lower than 35S:INTACT, so it is possible that this relates to RNA yield, although it was not the samples with the lowest yield that experienced failures and samples below the suggested RNA limit successfully generated libraries (60ng total RNA, while suggested limit for the kit was 100ng). If the accuracy of quantification at these lower RNA levels is noisy, it could be that the problem was indeed yield but that we cannot really know which samples are higher or lower yield. Due to the rarity of GCs, more plant material is required to obtain each KAT1:INTACT sample. It is therefore possible that there are some unknown secondary plant compounds being carried through and interferring with library preparation steps.

